# Effects of Ayahuasca on Ethanol-Conditioned Place Preference and ΔFosB Expression in the Nucleus Accumbens in Mice

**DOI:** 10.1101/2024.05.07.592962

**Authors:** Victor Distefano Wiltenburg, Gabriela Morales-Lima, Aline Valéria Sousa Santos, Marcela Echeverry Bermúdez, Fábio Cardoso Cruz, Gabriela de Oliveira Silveira, Maurício Yonamine, Paula Ayako Tiba, Fúlvio Rieli Mendes

## Abstract

**Background:** Ayahuasca, a psychoactive Amazonian preparation, is increasingly studied for substance-use disorders.

**Objectives:** Investigate whether oral lyophilised-ayahuasca attenuates ethanol-induced conditioned place preference (CPP) in mice and alters ΔFosB expression in nucleus accumbens (NAc).

**Methods:** Male Swiss mice received water or ayahuasca (130-1950 mg/kg, p.o.) 30 min before each of eight ethanol pairings (2g/kg i.p.) in a CPP paradigm. A separate cohort underwent acute toxicology (650–5000mg/kg) with behavioural-observation and rotarod. Alkaloids were quantified by LC-MS/MS. ΔFosB-immunoreactive nuclei were counted in NAc 24h after the CPP post-test.

**Results:** Alkaloids levels were within traditional ranges. High-dose ayahuasca(5000 mg/kg) produced transient-serotonergic-syndrome-like signs and rotarod locomotor-deficit; lower doses did not express toxicity. Ethanol produced a moderate-CPP in controls(ΔTime≈+60s), whereas ayahuasca-pretreatment abolished preference at all doses(ΔTime within±7s). One-way ANOVA on ΔTime showed a robust-Treatment effect(F(3,36)=8.83, p=0.00016); Tukey tests: control differed from each ayahuasca group (all p<0.05), with no differences among ayahuasca doses. ΔFosB density did not differ among groups(p>0.05).

**Conclusions:** Ayahuasca was well tolerated at ceremonies-equivalent doses and blocked ethanol-induced-CPP across all doses, while ΔFosB levels in NAc were unchanged at 24h. Limitations on the CPP baseline and ΔFosB results may limit sensitivity, generalisability and interpretation. Findings provide preliminary evidence that ayahuasca-pretreatment may blunt ethanol-context preference, reinforcing the need of replication with stronger reward baselines, naïve controls and complementary molecular markers.

## 1. Introduction

Alcohol-use disorder (AUD) remains a leading cause of preventable morbidity, yet available pharmacotherapies achieve only modest therapeutic effect (Bozkurt 2022). Psychedelic ethnomedicines such as Ayahuasca are emerging as a promising therapeutic option for conditions like substance use disorders (SUD) (dos Santos and Hallak 2020; Mendes et al. 2022; Wiltenburg et al. 2021). Ayahuasca is a psychedelic botanical decoction used by various traditional peoples in the Amazon for ritualistic and therapeutic purposes (McKenna and Riba 2016). It is usually prepared by boiling the bark and stems of the vine of *Banisteriopsis caapi* with the leaves of the shrub *Psychotria viridis*. The vine contains β-carboline alkaloids such as harmine, tetrahydroharmine (THH), and harmaline, while the shrub leaves contain N, N-dimethyltryptamine (DMT) (McKenna et al. 1984).

The β-carbolines inhibit gastric monoamine oxidase A (MAO-A), resulting in a synergistic effect that allows DMT to be orally active (McKenna and Riba 2016). DMT engages multiple 5-HT receptor subtypes (5-HT_2A_, _2C_, _1A_, _1B_, _1D, 7_), sigma-1 receptors and, at higher concentrations, trace-amine associated receptor 1 (Dodd et al. 2021; dos Santos and Hallak 2020; Grob et al. 1996; Nichols 2016; Speranza et al. 2017), producing a complex neurochemical profile with potential neuroplastic and anti-inflammatory effects.

Qualitative and quantitative observational studies support the therapeutic potential of ayahuasca on SUD treatment, where ceremony participants report lower prevalence of alcohol or cocaine dependence and reduced craving (Barbosa et al. 2018; Doering-Silveira et al. 2005; Jiménez-Garrido et al. 2020; Mendes et al. 2022; Thomas et al. 2013). Additionally, preclinical studies corroborate the potential of ayahuasca in drug addiction treatment (Daldegan-Bueno et al. 2023; Mendes et al. 2022). Ayahuasca administration has been shown to reduce voluntary amphetamine consumption, amphetamine-induced locomotor activity and anxiety in rats (Godinho et al. 2017), attenuate naloxone-induced withdrawal syndrome in morphine-dependent rats (Aricioglu-Kartal et al. 2003), and inhibited ethanol-induced sensitization and reduced the ethanol withdrawal symptoms in mice (Almeida et al. 2021; Oliveira-Lima et al. 2015). Collectively, these studies indicate that ayahuasca interacts with multiple drug classes, yet its effect on the acquisition of ethanol-context associations remains unknown.

Conditioned place preference (CPP) has been used to investigate SUD and provides a pragmatic translational assay for drug-context learning. Two studies showed that ayahuasca suppressed the expression of ethanol CPP when administered after conditioning (Cata-Preta et al. 2018; Gianfratti et al. 2022). Whether pre-treatment can prevent CPP acquisition, and how this relates to ΔFosB, remains unknown. Addressing this gap is essential, because CPP can be sensitive to interventions that affect mesolimbic reward circuitry and can help identify anti-addictive potential (Daldegan-Bueno et al. 2023; Mendes et al. 2022).

Additionally, repeated drug exposure has been shown to induce the stable transcription factor ΔFosB in D1-medium spiny neurons of the nucleus accumbens (NAc), a change that can promote behavioural sensitisation and increase relapse risk. Ethanol elevates ΔFosB in both NAc core and shell, and the reduction of this signal is associated with reduced drug-seeking behaviour (Robison and Nestler 2022; Wille-Bille et al. 2021).

There is a need for novel interventions that target the mesocorticolimbic circuitry encompassing the ventral tegmental area, NAc and orbitofrontal cortex (Bracht et al. 2021). Building on ayahuasca’s emerging anti-addictive profile, the present study investigated whether pretreatment with lyophilised ayahuasca using doses spanning sub- to supra-ritual human equivalents to maximise translational relevance, would attenuate ethanol-induced conditioned place preference (CPP) and modify ethanol-related ΔFosB accumulation in the NAc.

## 2. Materials and Methods

### 2.1. Botanic material

The ayahuasca preparation was provided by the Scientific Commission of the União do Vegetal (UDV). Samples of the plants used to produce the brew were collected in the city of Mogi das Cruzes, São Paulo State, Brazil (−23.5228, −46.1883), and identified as *Banisteriopsis caapi* (Spruce ex Griseb.) Morton (Malpighiaceae) and *Psychotria viridis* Ruiz & Pav (Rubiaceae).

The voucher specimens were deposited in the Herbarium of the Federal University of ABC (UFABC) under registration numbers HUFABC002262 *(B. Caapi)* and HUFABC002263 *(P. Viridis*). The collection and research permits complied with Brazilian biodiversity law (Sisgen #AF827A4)

### 2.2. Ayahuasca preparation and lyophilization

The ayahuasca preparation was concentrated (to approximately 25% of original volume) under reduced pressure (rotary evaporator, 45 °C, in vaccum), frozen at −20 °C and lyophilized (Liotop k105) to produce a dry extract. The brew concentration was determined as 356 mg/ml. The powder was stored in amber screw-cap vials in vacuum desiccator at room temperature and resuspended in distilled water (pH 7.2) as needed.

### 2.3. Quantification of alkaloids

A lyophilized ayahuasca sample was resuspended in distilled water to achieve the original concentration and the β-carbolines (harmine, harmaline, and THH) and DMT were quantified by high-performance liquid chromatography (HPLC) coupled with a mass spectrometer (LC-MS /MS), according to the published method of in de Oliveira Silveira et al. (2020). An HPLC profile (chromatogram) of the sample is located in the supplementary material Figure S1.

### 2.4. Animals

Male Swiss mice (2 to 3 months old, sample size specified in sessions 2.6 and 2.7) were obtained from the animal facilities of Federal University of São Paulo (CEDEME/UNIFESP) and Federal University of ABC (UFABC Bioterium). The mice were housed in micro-insulated polypropylene cages (30 × 19 × 13 cm, 5 animals per cage, with minimal enriched environment, such as paper nestlets) under a 12h/12h light/dark cycle (lights off at 7:00 pm) and controlled temperature (21 ± 4 ◦C). Water and food (Nuvilab^®^) were provided *ad libitum*. The mice were habituated for 15 days prior to the start of the experiments, conducted during the light phase. Mice were randomly assigned to treatment groups using a random-number generator and experimenters were blinded to treatment during Toxicological assessment. All procedures adhered to Brazilian CONCEA guidelines and ARRIVE 2.0 recommendations and were approved by the UFABC Animal Ethics Committee (protocol 2502090217).

### 2.5. Drugs and treatments

Lyophilized ayahuasca (dry extract) was diluted in distilled water and administered by gavage (orally, v.o.) at a volume of 10 ml/kg weight. The doses used for all behavioral and pharmacological tests were calculated based on a human adult (∼70kg) average consumption of ayahuasca at UDV members during a routine religious ceremony (128ml), which is equivalent to a dose of approximately 650 mg/kg. Doses of 130 (0.2X unitary), 650 (1X unitary), 1950 (3X unitary), 3560 (5X unitary) and 5000 mg/kg (maximum recommended dose in toxicological studies) were used. Control animals received vehicle gavage (water, 10 ml/kg) and saline i.p. when needed for experimental models, and specific doses used in each experiment are addressed in the sections 2.6 and 2.7). Absolute ethanol (Synth, Brazil) was diluted in saline solution (0.9% sodium chloride) at a concentration of 20% and injected in a volume of 14 ml/kg to achieve a dose of 2 g/kg i.p.) route.

### 2.6. Toxicological assessment – Experiment 1

A separate cohort of 28 male Swiss mice were divided into four groups (n = 7/group) and received an oral dose of water (control) or increasing doses of ayahuasca (650, 3560, and 5000 mg/kg). Animals were placed in stainless-steel wire cages immediately after administration and observed by a blinded experimenter at intervals of at 0–15, 15–30, 30–60, 60–120, 120–240 minutes and 24 hour. Presence or absence of several signs and behaviors (such as signs of stereotypy, Straub’s tail, in addition to trembling and ruffled fur) were registered, according to (Carlini 2011).

After the screening, animals motor coordination were evaluated on a constant-speed rotarod (t-off 180 seconds). Each mouse performed three trials at baseline, then at 45, 90 and 180 min after ayahuasca administration. The latency to first fall and number of falls were averaged per time-point, as described in (Carlini 2011). Data were analyzed by two-way repeated-measures ANOVA (Dose × Time) as detailed in Section 2.10.

### 2.7. Conditioned place preference (CPP) - Experiment 2

The CPP apparatus consisted in two distinct chambers (20 × 20 × 40 cm) separated by a neutral corridor (12 × 12 × 40 cm) and differed in wall color (black vs white), floor texture (grid vs mesh) and visual cues. After each session the chambers were cleaned with 70% ethanol and waited one hour before the next usage.

The time mice spent in each chamber was recorded and analyzed using ToxTrac software (https://toxtrac.sourceforge.io). The procedure (adapted from Prus et al. (2009) consisted of two habituation sessions (one session/day), one 15-min pre-conditioning test, eight daily conditioning sessions (four ethanol, four saline, alternated), and a 15-min post-conditioning test 24 h after the last conditioning session, without any treatment (Figure 1).

**Figure 1.**
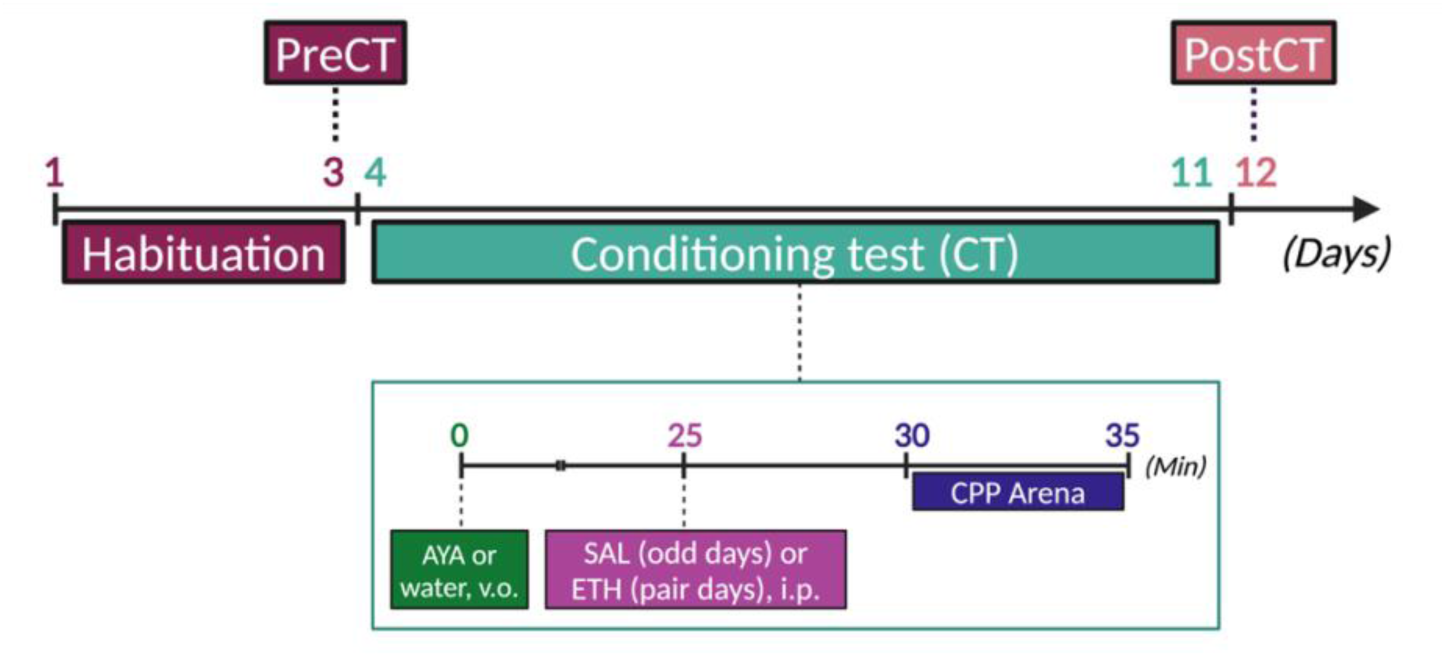
Experimental timeline for ethanol-conditioned place preference (CPP) and ayahuasca pretreatment. The CPP apparatus consisted of two chambers with different characteristics, such as colour, floor texture and visual stimuli, and a neutral corridor connecting the two chambers. Habituation - mice freely explored both chambers for 15 minutes (for two days). Pre-conditioning test (PreCT) - drug-free 15 min test to assess baseline chamber preference at day 3. Conditioning - mice were divided into four experimental groups: Water (control), Ayahuasca (AYA) 130, 650 or 1950 mg/kg. The conditioning sessions were carried out over 8 days. Ayahuasca or water was administered orally 30 minutes before each session (time = 0). Ethanol (ETH, 2.0 g/kg) or saline (SAL) was administered 5 minutes before the animals were confined to a chamber (time = 25) for 5 minutes (four days each and alternating). Drug-paired chamber assignment was counter-balanced within each group. Post-conditioning test (PostCT), 24 h later, drug-free mice freely explored both chambers for 15 min. Position was tracked automatically (ToxTrac).

#### 2.7.1. Habituation and preconditioning test

Mice were placed in the neutral corridor with both doors open for three consecutive days (15 min day⁻¹) to habituate to the apparatus. On day 3, a 15-min pre-conditioning test established baseline preference. The time animals spent in each chamber were recorded and analyzed with ToxTrac. Animals displaying a strong side bias (>70% time in one chamber) were re-assigned to balance groups (none exceeded exclusion criteria). No injections were administered on either the habituation days or the pre-conditioning test.

#### 2.7.2. Conditioning

After the pre-conditioning test, animals were evenly distributed among four experimental groups (n = 9/10, total animals = 39): control water, ayahuasca 130, 650, and 1950 mg/kg. To reduce possible bias, half of the mice were paired with ethanol in the less preferred chamber, and the other half in the more preferred chamber in the pre-conditioning test (Prus et al. 2009).

The conditioning sessions were performed for eight consecutive days. Animals received oral ayahuasca or water (10 ml/kg). **2**5min later they were injected i.p. with ethanol (2 g/kg, 20% v/v) or saline. Five minutes after injection they were confined to the appropriate chamber for 5 min. Ethanol and saline days alternated (E–S–E–S–E–S–E–S. Described in Figure 1).

#### 2.7.3. Postconditioning test

Twenty-four hours after the last conditioning session, mice were returned to the apparatus with doors open for a 15-min post-test. Time in the ethanol-paired and saline-paired chamber (primary outcome) was recorded and later analyzed (Section 2.11). Researchers scoring behavior and conducting histology were blinded to group. At 24 h post-test, mice were deeply anaesthetized and transcardially perfused with phosphate-buffered saline followed by 4% paraformaldehyde. Brains were post-fixed overnight and processed for ΔFosB immunohistochemistry (Section 2.8).

### 2.8. Tissue Preparation

Twenty-four hours after the CPP post-test, mice were deeply anaesthetized with urethane (1.5 g/kg, 30% w/v, i.p. Sigma-Aldrich, St. Louis, MO, USA) and transcardially perfused with ice-cold phosphate-buffered saline (PBS, 0.1 M, pH 7.4; 30 mL) followed by 4% paraformaldehyde (PFA, Sigma-Aldrich, St. Louis, MO, USA) in PBS (40 mL). Brains were removed, post-fixed in 4% PFA overnight at 4 °C, rinsed in PBS and cryoprotected in 30% sucrose/PBS for 48h at 4 °C. Tissues were flash-frozen on dry ice and stored at –80 °C until sectioning. Coronal sections (30 µm) were cut on a cryostat (Leica CM1860 UV Cryostat; Leica Microsystems Inc, IL, USA) at −15° to −20°C. Serial sections taken through the striatum (containing the NAc) were collected in ethylene glycol anti-freezing solution and stored at −20°C until the day of the immunohistochemistry assay(Prieto et al. 2023) at –18 °C and collected serially into ethylene glycol anti-freezing solution at –20 °C. Every sixth section containing the nucleus accumbens (NAc; ∼1.70 mm to 0.62 mm anterior to bregma) was reserved for ΔFosB immunohistochemistry (see Section 2.9). All tissue processing and subsequent image analyses were performed by investigators blinded to treatment assignment (Prieto et al. 2023).

### 2.9. Free-floating immunohistochemistry

Coronal sections containing nucleus accumbens were washed in a 0.1 M phosphate buffer solution (PBS) and incubated with 10 mM sodium citrate buffer at 70 °C for 20-30 minutes. The sections rested at room temperature for 30 minutes and were then washed with 0.1M PBS with Triton X-100 (PBST) and blocked with 2% bovine serum albumin (Sigma-Aldrich Brazil, São Paulo, Brazil) in PBST. The tissues were incubated overnight at 4°C with anti-ΔFosB primary antibody (ΔFOSB; rabbit; Cell Signaling Technology, Boston, US, ref. 2251; 1:1000 dilution) in blocking solution. Then, sections were washed in PBST and incubated for 2-3 hours with biotinylated anti-rabbit secondary antibody (1:450, Vector Laboratories, Burlingame, CA, USA). Finally, the sections were washed in PBST and incubated for 90-120 min in the avidin-biotin-peroxidase complex (ABC Elite kit, PK-610, Vector Laboratories) in PBST, washed and then incubated with 3,3′-diaminobenzidine for approximately 5 min. The sections were transferred into 0.1M PBS and mounted onto chrome alum-gelatin-coated slides. Once dry, the sections were dehydrated with a graded series of ethanol (distilled water; ethanol 30%, 60%, 90%, 95%, and 100%) and xylol before cover slipping with Entellan (Merck, Darmstadt, Germany). A sample of negative control with absence of primary antibody was always carried out to show the absence of non-specific binding.

### 2.10. Image analysis

ΔFosB-immunoreactive nuclei were quantified in the nucleus accumbens (NAc) core and shell between ∼1.70 mm and ∼0.62 mm relative to bregma (Franklin and Paxinos 2013). Every sixth section produced 3–6 bilateral images per animal. Images were captured under fixed illumination with a Leica DM5500 microscope and DFC295 camera at 1.25x/0.0375 and 5x/0.12 objectives. ΔFosB positive cells were quantified using ImageJ software (https://imagej.nih.gov/), measured by the density of positive cells per 0.1 mm². Density data were analysed by one-way ANOVA as detailed in Section 2.11.

### 2.11. Statistical analysis

The data was analyzed using JAMOVI v2.4.0 (https://www.jamovi.org/) and GraphPad Prism v7 (GraphPad Software, CA, USA). Normality was assessed by Shapiro–Wilk and inspection of Q–Q plots. Homogeneity of variance by Levene’s test and, for repeated-measures designs, sphericity by Mauchly’s test. No outliers (>3 SD from the mean) were detected or removed.

Toxicological assessment behaviour was qualitatively analysed and the rotarod latency compared across Dose and Time by two-way repeated-measures ANOVA. Conditioned place preference (CPP), time in the ethanol or saline-paired chamber, was analysed with a 4 (Treatment) × 2 (Session: Pre vs Post) repeated-measures ANOVA, one way ANOVA (Treatment) and pre vs post t-test. ΔFosB-positive cell density in NAc core and shell was compared across the four treatment groups with one-way ANOVA (factor: Treatment). When an omnibus F-test was significant (α = 0.05, two-tailed), Tukey’s was applied for pairwise comparisons.

## 3. Results

### 3.1. Quantification of alkaloids

The quantification of alkaloids (LC-MS/MS) showed the concentrations in the used ayahuasca as (mean ± SD, n = 3): harmine 1.4 ± 0.05 mg/ml, harmaline 0.18 ± 0.01 mg/ml, THH 1.86 ± 0.04 mg/ml and DMT 0.08 ± 0.02 mg/ml. These values fall within the ranges reported in the literature, although DMT concentration is relatively low (Callaway et al. 2005). At the behavioural unitary dose (650 mg/kg), mice therefore received approximately 0.15 mg/kg DMT, 2.56 mg/kg harmine, 0.33 mg/kg harmaline and 3.4 mg/kg THH. Intra- and inter-day relative standard deviations were below 8%, and all analytes were above their limits of quantification.

### 3.2. Toxicological assessment

No mortality occurred during the toxicological assessment and only the 5000 mg/kg group displayed signs of stereotypy, Straub’s tail, tremor and pilo-erection during the first 60 min, which declined substantially in the following hour, and were completely absent by the 2–4 h interval and at 24 h. Neither 650 mg/kg nor 3 560 mg/kg produced any observable signs at any time-point (0/7 animals for every behaviour).

Rotarod performance showed a significant Dose × Time interaction (two-way RM-ANOVA, F (12, 72) = 4.11, p < 0.001; Figure 2). Post-hoc Tukey tests revealed that mice receiving 5 000 mg/kg exhibited a transient reduction in latency to fall (Rota-rod performance) at 45 min (mean ± SEM = 14.1 ± 3.2 s) compared with both vehicle controls (53 ± 4.4 s, p = 0.008) and their own baseline (p < 0.001). Performance at 90 min and 180 min were indistinguishable from control (both p > 0.10). Performance values for the 130 mg/kg and 3,560 mg/kg groups did not differ from control at any time-point (all p > 0.10).

**Figure 2.**
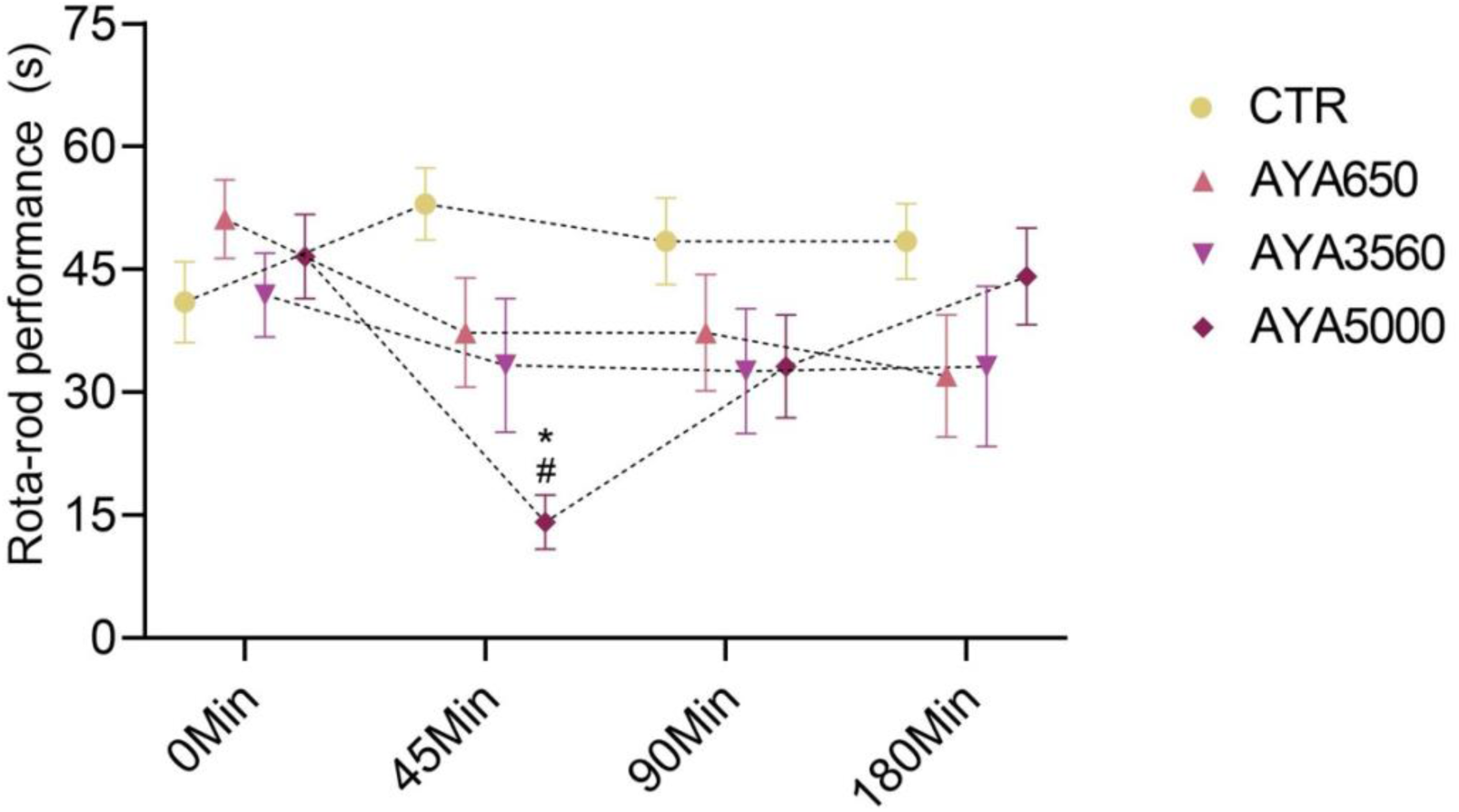
Rotarod performance following acute oral ayahuasca in the toxicological assessment. Latency to fall (Rota-rod Performance, mean ± SEM) on a constant-speed rotarod (12 rpm) at baseline (0) and 45, 90, 180 min after dosing with water (CTR) or ayahuasca (AYA 650, 3 560, 5 000 mg/kg). Two-way RM-ANOVA (Dose × Time) showed a significant interaction (*F*(12,72)=4.11, *p*<0.001). The 5 000 mg/kg group exhibited a transient deficit at 45 min vs. Control and its own baseline (Tukey’s post hoc, *p*<0.05); performance normalised by 90–180 min. No effects were detected for 650 or 3 560 mg/kg at any time point. *n=7 per group*.

### 3.3. Conditioned place preference

Baseline exploration did not differ among groups: a 4 (Treatment) × 2 (Chamber) repeated-measures ANOVA on pre-test times showed no main effects or interaction (*F*(3,35)=0.73, *p*=0.55). Behavioral changes were quantified by measuring the time spent in the ethanol-paired and saline-paired chambers (here named non-paired). ΔTime was calculated by subtracting the time spent in the non-paired chamber from the time spent in the ethanol-paired chamber (Fig. 3A and 3B). A one-way ANOVA revealed a significant Treatment effect on ΔTime (*F*(3, 36)=8.83, *p*=0.00016, partial η² = 0.42). Control mice displayed robust CPP (+60 ± 0.73 s), whereas ΔTime was near zero in every ayahuasca group (AYA 130: +6 ± 7.27 s; AYA 650: +7 ± 8.54 s; AYA 1950: +5 ± 14.23 s). Tukey post-hoc tests (critical *q*=3.77; HSD=34.3 s) showed the Control group differed from each ayahuasca dose (mean differences 53–55 s, *q* ≥ 5.8, *p* < 0.001), while the ayahuasca groups did not differ among themselves (|Δ| ≤ 2 s, *q* ≤ 0.22, *p* > 0.90). One-sample *t*-tests confirmed that ΔTime did not differ in any ayahuasca group (all *p* > 0.15), indicating absence of CPP, whereas Control mice showed a strong preference *p*<0.021. Thus, despite analytic limitations, there is a clear separation between ethanol-only controls and all ayahuasca-pretreated groups, with ayahuasca blocking ethanol-induced CPP across the entire dose range.

**Figure 3.**
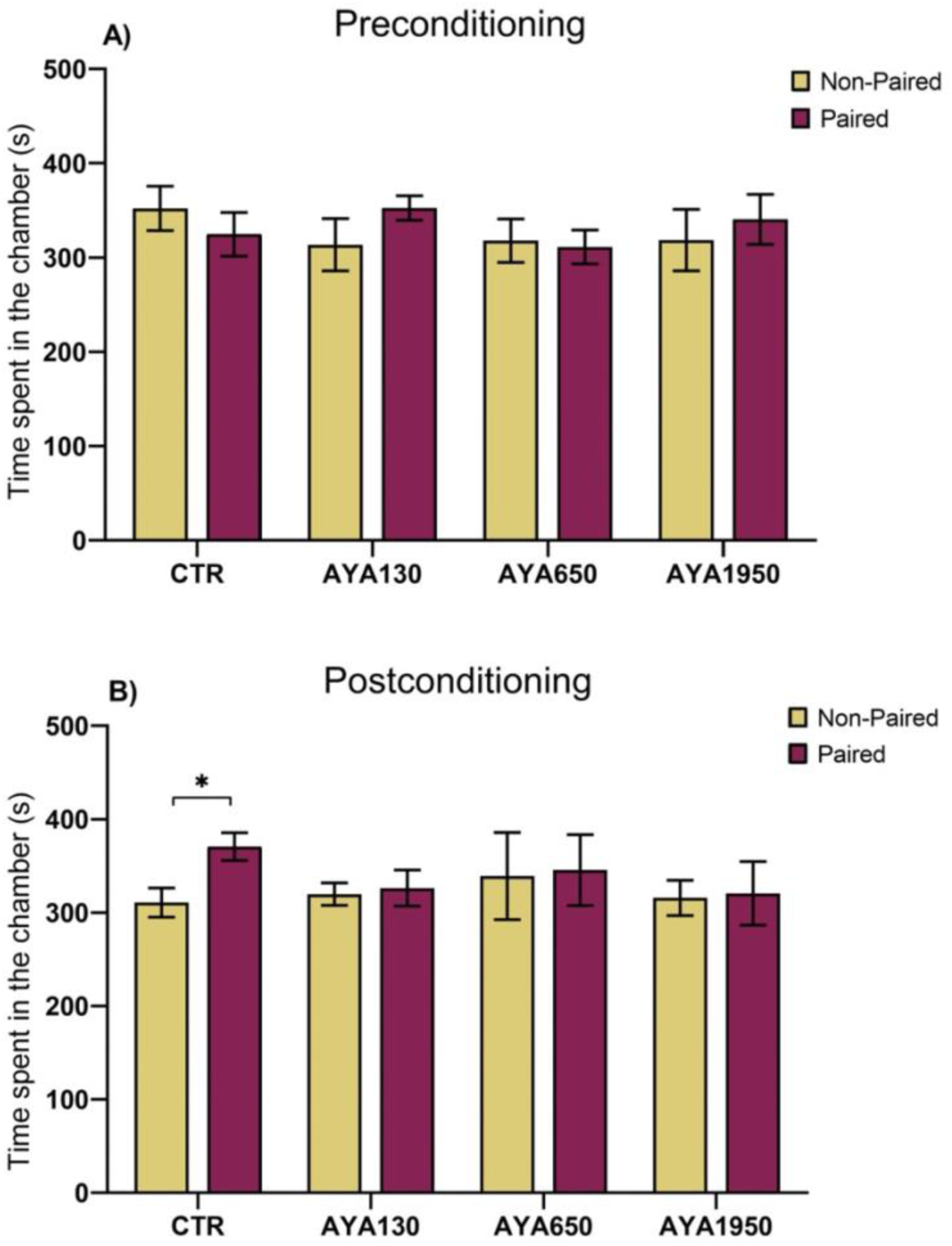
Ayahuasca pretreatment and ethanol-induced conditioned place preference (CPP). Time spent (seconds) in saline-paired (non-paired) and ethanol-paired (paired) chambers by mice administered orally with water (CTR), or ayahuasca (AYA) 130, 650, and 1950 mg/kg, 25 minutes before intraperitoneal administration of 0.9% saline (non-paired) and 2.0 g kg⁻¹ ethanol (paired). A) Pre-conditioningtest (PreCT) time (seconds) in each chamber (mean ± SEM) shows no baseline chamber preference across groups (4 Treatment × 2 Chamber RM-ANOVA, *F*(3,35)=0.73, *p*=0.55). B) Post-test (PostCT) time in chambers. Change score ΔTime = (post paired) − (post non-paired) per group (mean ± SEM). One-way ANOVA on ΔTime: *F*(3,36)=8.83, *p*=0.00016, partial η²=0.42. Tukey HSD: Control differs from each AYA dose (*p<0.05), AYA doses do not differ among themselves (p>0.15). Control shows robust CPP (+60 ± 0.73 s), whereas ΔTime ≈ 0 in all AYA groups (AYA 130: +6 ± 7.27 s; AYA 650: +7 ± 8.54 s; AYA 1950: +5 ± 14.23 s). n=9–10 per group.

### 3.4. ΔFosB protein immunoreactivity

Immunoreactive nuclei were quantified in the shell and core of the NAc (Fig. 4A,B). A one-way ANOVA revealed no main effect of Treatment on ΔFosB-positive cell density in either region (core: *F*(3, 36)= 2.17, p = 0.243; shell: *F*(3, 36)= 3.83, p = 0.077; Fig. 4C,D).

**Figure 4.**
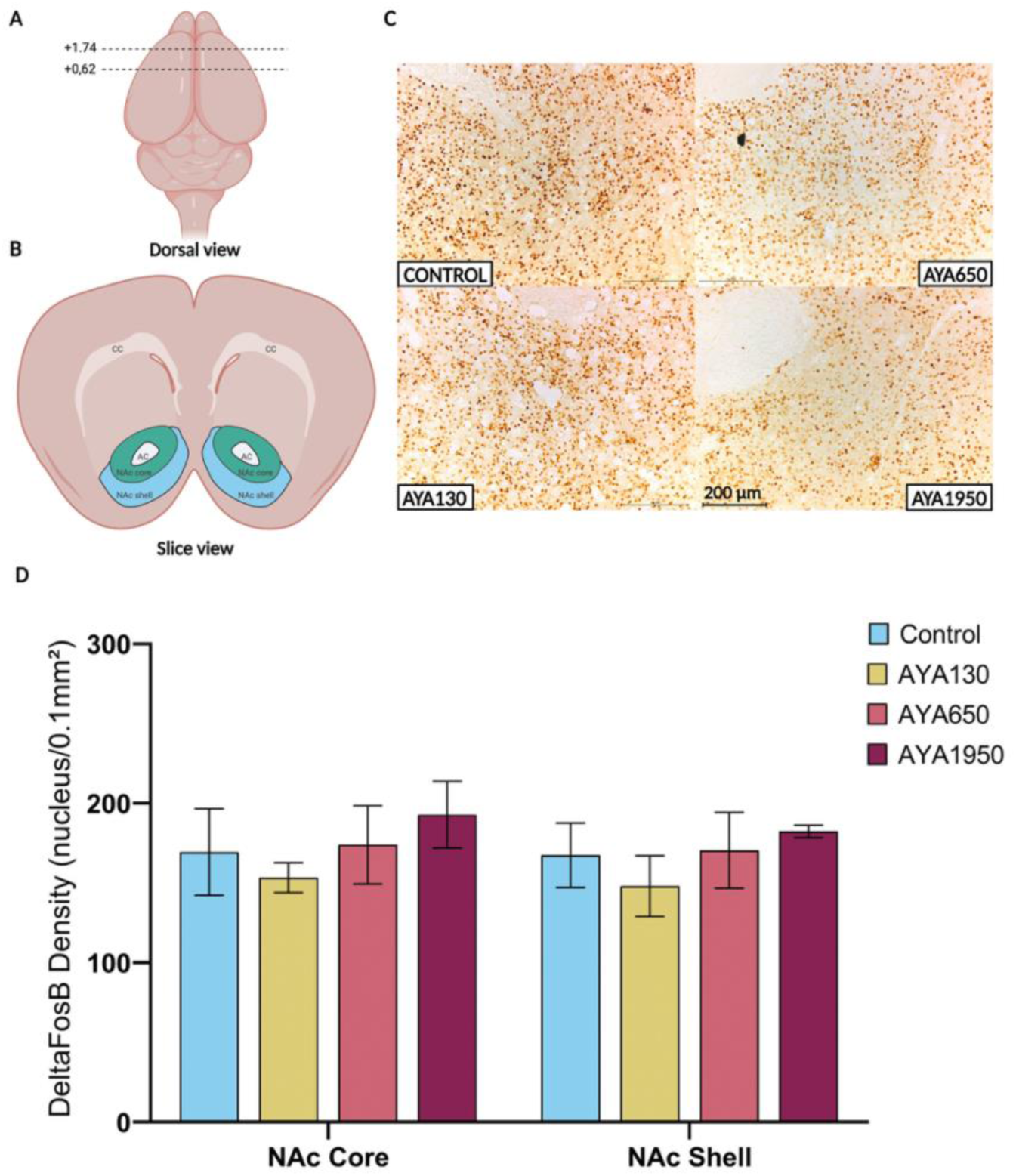
ΔFosB immunoreactivity in nucleus accumbens (NAc) after CPP. A) Dorsal schematic showing the rostro-caudal range of coronal sections analysed (approx. +1.70 to +0.60 mm from bregma). B) Example section indicating NAc core and shell (AC, anterior commissure; CC, corpus callosum). C) Representative photomicrographs of ΔFosB-immunoreactive nuclei in core and shell (Cell Signaling #22509S; DAB). Scale bars: 200 µm. D) Quantification of ΔFosB-positive nuclei (nucleus per 0.1 mm²; mean ± SEM) in Control and AYA 130, 650, 1950 mg/kg groups after CPP protocol. One-way ANOVA showed no Treatment effect in either subregion (*p*>0.05). n=4–5 brains per group; sections and counting performed to treatment.

## 4. Discussion

Several observational studies link ceremonial ayahuasca use to lower prevalence of alcohol and cocaine dependence and improved psychological wellbeing (Mendes et al. 2022; Wiltenburg et al. 2021). Pre-clinical research also reports anti-addictive effects, but most studies rely on heterogeneous preparations and lack integrate behavioural and molecular investigation (Daldegan-Bueno et al. 2023; Gianfratti et al. 2022; Nunes et al. 2016). By combining phytochemical analysis, acute toxicology, ethanol-conditioned place preference (CPP) and ΔFosB immunohistochemistry, the present study advances that evidence base.

The phytochemical analysis of the ayahuasca used in this study revealed β-carbolines at literature-consistent concentrations, while DMT levels (0.08 mg/ml) were below the averages typically reported but still within the documented range (Callaway 2005; Souza et al. 2019). Notably, the ayahuasca sample induced the desired subjective effects during the group’s ritual, as confirmed by UDV members (personal communication). The doses used in this study were based on the average ritualistic unitary dose calculated for an individual of 70 kg, which is 651 mg/kg (as shown on session 2.5).

In the toxicological assessment, oral ayahuasca up to 1 950 mg/kg (3 × unitary dose) produced no observable signs or motor deficit, matching previous reports of low toxicity in rodents, or any other behavioural impairment (Morais 2014). The highest dose (5 000 mg/kg) showed transient serotonergic-syndrome-like behaviours (Carlini 2011) and rotarod impairment that vanished within four hours. There were no fatalities. These findings reinforce the relative safety of ritual-equivalent dosing (Morais 2014; Pires et al. 2010).

In the conditioned place preference experiment, baseline exploration was equivalent across groups (pre-test RM-ANOVA was not significant), so post-test differences reflect treatment. The control group (ethanol only) developed a moderate preference (ΔTime ≈ +60 s), whereas ayahuasca pretreatment blocked preference at all three doses (ΔTime within ±7 s). A one-way ANOVA on ΔTime confirmed a robust treatment effect (F(3, 36) = 8.83, p = 0.00016, partial η² = 0.42), and Tukey post hoc tests revealed clear separation between the control group and each ayahuasca dose (all p < 0.05), with no differences among the ayahuasca doses. In practical terms, despite the limitation on the moderate baseline CPP, ayahuasca pretreatment reduced the mean preference score relative to the control group, indicating robust and clear attenuation of ethanol-CPP across the tested dose range. Although the ANOVA of ΔTime was performed using group summaries rather than a full repeated-measures model on individual pairs, the convergence of the omnibus test, post hoc contrasts, and one-sample tests (ΔTime ≈ 0 in all ayahuasca groups) supports the conclusion that ayahuasca blocks ethanol-induced CPP.

Previous studies have also observed that ayahuasca blocks ethanol-induced CPP at different doses and that ayahuasca might induce CPP by itself (Cata-Preta et al. 2018; Gianfratti et al. 2022). On the other hand, Cata-Preta et al. (2018) showed that the single extracts of *P. viridis* and *B. caapi* did not induce CPP *per se*, nor blocked ethanol-induced CPP, highlighting the importance of the synergistic effects of combining ayahuasca constituents. Furthermore, a 2024 study suggests that the preference of ayahuasca over water in the two-bottle choice procedure depends on the frequency of exposure and the concentration of ayahuasca (Kisaki et al. 2024).

ΔFosB is known to be a transcription factor that accumulates in the brain after any drug exposure. It is a stable molecule that can remain present for long periods of time (Damez-Werno et al. 2012; Nestler et al. 2001). ΔFosB is also known to increase the sensitivity of the reward circuit to a drug and may be one of the mechanisms behind drug-induced craving and euphoria (Nestler et al. 2001). Interestingly, structural changes in the NAc caused by ΔFosB are also considered to cause drug relapse, and the overexpression of the protein is known to occur in D1 neurons of the NAc of people suffering from AUD (MacNicol 2017; Olsen 2011). This mechanism through ΔFosB may produce sensitivity in former addicts when they are reintroduced to drugs, leading to rapidly escalating relapse (MacNicol 2017).

To our knowledge, this is the first study to evaluate ΔFosB-positive cells after ayahuasca treatment. Despite behavioural attenuation, there were no differences in the number of ΔFosB-positive nuclei in the NAc core and shell among groups (core: F(3, 36) = 2.17, p = 0.243; shell: F(3, 36) = 3.83, p = 0.077). This lack of the change in ΔFosB-positive nuclei may be explained by the lack of statistical power, since our design had 80% power to detect a ≥35% change, so smaller shifts could go undetected, and timing, since ΔFosB peaks after repeated or prolonged drug exposure (Damez-Werno et al. 2012), whereas we evaluated 24 h post-test. Future work should include a naïve, no ethanol-ayahuasca and ayahuasca-only group for CPP and ΔFosB.

To understand the hypothesis of how ayahuasca can be used therapeutically for AUD, it is important to emphasize that addiction is known to be a chronic disease that can impair the ability of the brain’s reward circuitry to respond to reward, as well as the modulation of stress and self-regulation (Volkow et al. 2019). Addictive drugs have the potential to increase dopamine release in the mesolimbic reward pathway, either directly by acting on the dopaminergic system or indirectly by affecting the modulatory activity of other neurotransmitters, such as serotonin (Tomkins and Sellers 2001; Volkow et al. 2019).

On a mechanistic note, β-Carbolines (harmine, harmaline, and to a lesser extent, tetrahydroharmine THH) inhibit MAO-A, which elevates synaptic serotonin (5-HT). By acting on 5-HT_2C_ receptors (and, to a lesser extent, 5-HT_2A_ receptors), they can dampen mesolimbic dopamine output. This occurs, in part, via the excitation of GABAergic interneurons that inhibit dopaminergic neurons (Howell and Cunningham 2015; Sari et al. 2011; Sullivan et al. 2015). THH’s weak SERT inhibition may reinforce this 5-HT2C effect. Given the low DMT content of ayahuasca, a β-carboline-driven serotonergic mechanism rather than strong 5-HT2A-mediated psychedelic signalling is a possible explanation for the anti-CPP effect. However, receptor specificity remains hypothetical and warrants antagonist and constituent isolation studies.

Observational studies emphasise set-and-setting as therapeutic co-factors (Liester and Prickett 2012; Mendes et al. 2022). However, our behaviour-based protocol excluded these psychosocial elements, yet ayahuasca still attenuated ethanol CPP, suggesting an intrinsic pharmacological component. Nonetheless, behaviour tests may not be sufficient to investigate the totality of the potential effect of ayahuasca in drug addiction. Clinical translations are important to further the discussion and will require placebo-controlled trials paired in supportive settings akin to modern psychedelic-assisted therapy.

To summarize, within ceremony-equivalent doses, lyophilised ayahuasca was acutely safe, preserved motor coordination, and blocked ethanol-induced CPP without detectable changes in ΔFosB in the NAc at 24 h. These preclinical data extend evidence for anti-addictive potential and provide a preliminary foundation for mechanistic studies and controlled clinical testing in alcohol-use disorder. Further studies that address this paper’s limitations and expand to additional paradigms are needed to clarify mechanisms and validate ayahuasca’s impact on alcohol-related reward.

## Data availability statement

The data that support the findings of this study are available from the corresponding author, VDW, upon reasonable request.

---

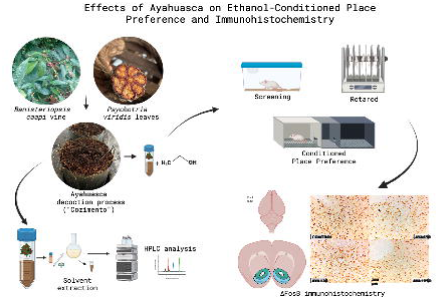

