## Supplemental Figure S1 for "Effects of Ayahuasca on Ethanol-Conditioned Place Preference and ΔFosB Expression in the Nucleus Accumbens in Mice"

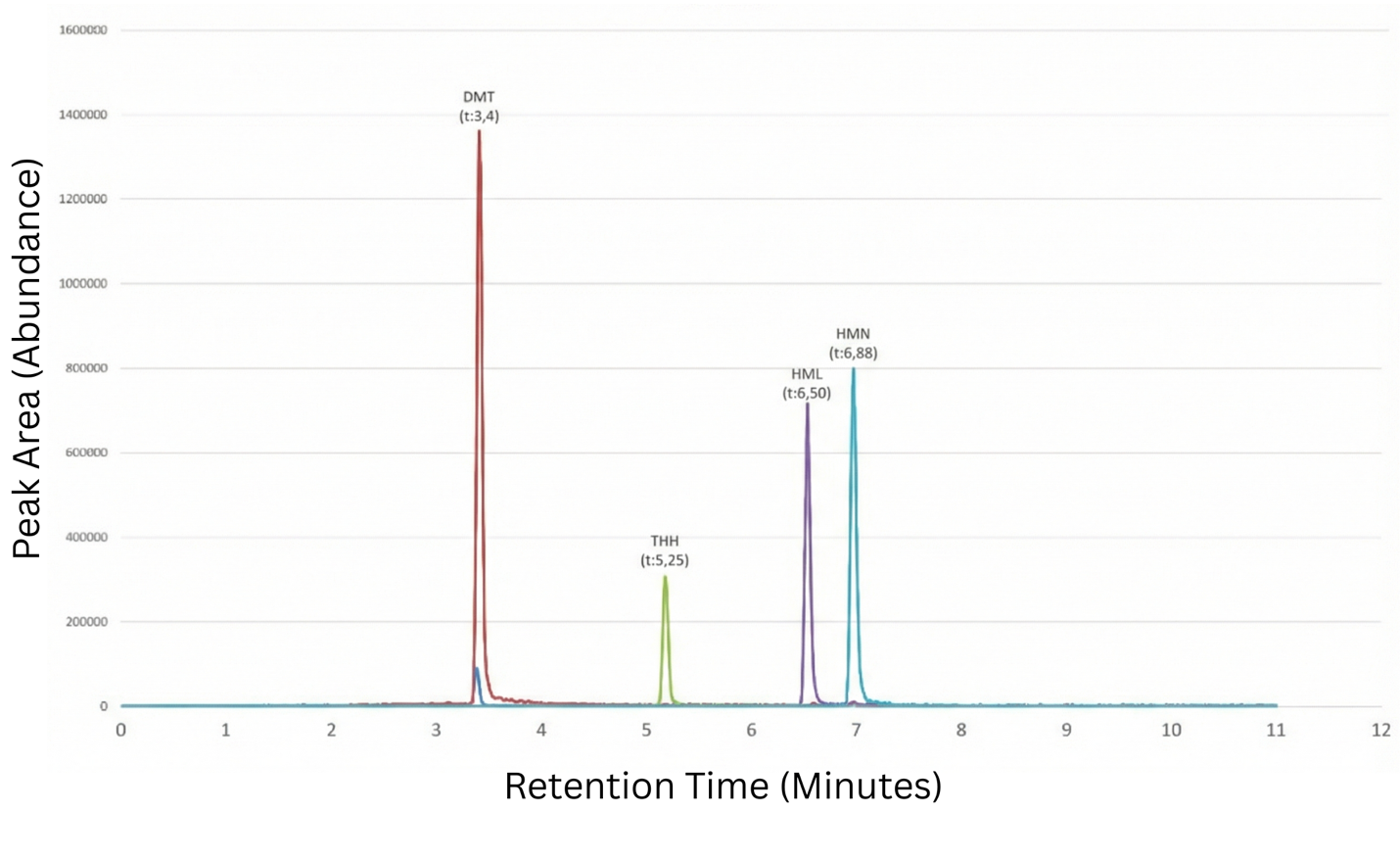

**Figure S1.** LC-MS chromatogram obtained from a sample containing 1. dimethyltryptamine (DMT), 2. DMT-*d*_6_ (internal standard), 3. tetrahydroharmine (THH), 4. harmaline (HRL) and 5. harmine (HRM) after the dilute-and-shoot procedure. The alkaloids were quantified by high-performance liquid chromatography (HPLC) coupled with a mass spectrometer (LC-MS /MS), according to the published method of in de Oliveira Silveira et al. (2020).
